# Cortical Dynein Drives Centrosome Clustering in Cells with Centrosome Amplification

**DOI:** 10.1101/2022.08.04.502862

**Authors:** Dayna L. Mercadante, William A. Aaron, Sarah D. Olson, Amity L. Manning

## Abstract

During cell division, the microtubule nucleating and organizing organelle, known as the centrosome, is critical for the formation of the mitotic spindle. In cells with two centrosomes, each centrosome functions as an anchor point for microtubules, leading to the formation of a bipolar spindle and progression through a bipolar cell division. When extra centrosomes are present, multipolar spindles form and the parent cell may divide into more than two daughter cells. Cells that are born from multipolar divisions are not viable and hence clustering of extra centrosomes and progression to a bipolar division are critical determinants of viability in cells with extra centrosomes. We combine experimental approaches with computational modeling to define a role for cortical dynein in centrosome clustering. We show that centrosome clustering fails and multipolar spindles dominate when cortical dynein distribution or activity is experimentally perturbed. Our simulations further reveal that centrosome clustering is sensitive to the distribution of dynein on the cortex. Together, these results indicate that dynein’s cortical localization alone is insufficient for effective centrosome clustering and instead, dynamic relocalization of dynein from one side of the cell to the other throughout mitosis promotes timely clustering and bipolar cell division in cells with extra centrosomes.

## Introduction

Proper formation and maintenance of the dynamic mitotic spindle is necessary to ensure accurate segregation of chromosomes and cell division resulting in two viable daughter cells. Assembly of the spindle is initiated by the nucleation of microtubules at each of two centrosomes, the organelles that function as the dominant microtubule organization center of the cell (Hinchliffe, 2011; Petry, 2016). Microtubule plus-ends radiate outward from the centrosome and interact with the cell boundary, other microtubules, and chromosomes (Forth and Kapoor, 2017). In response to mechanical forces on and by the microtubules, the centrosomes are positioned and ultimately become the two poles of a bipolar spindle (di Pietro et al., 2016; Busson et al., 1998; Goshima et al., 2005; Kiyomitsu and Cheeseman, 2012; Merdes et al., 2000; Okumura et al., 2018).

Dynein is the major minus-end directed motor protein in the cell and has many functions. Throughout mitosis, dynein is localized at the kinetochore on chromosomes where it contributes to chromosome movement and alignment (Bader and Vaughan, 2010), at the spindle-poles where it actively sustains focusing of microtubule minus ends at centrosomes (Merdes et al., 2000), and at the cell cortex, where it contributes to centrosome movement and thus impacts bipolar spindle length, orientation, and positioning (Busson et al., 1998; O’Connell and Wang, 2000; Kiyomitsu and Cheeseman, 2012; di Pietro et al., 2016; Okumura et al., 2018; Mercadante et al., 2021). Dynein and its binding partner dynactin are anchored at the cell cortex by a complex including the proteins LGN, Afadin, and NuMA (Du and Macara, 2004; Kiyomitsu and Cheeseman, 2012; Kotak et al., 2012, 2013; di Pietro et al., 2016; Okumura et al., 2018). Cortical localization of dynein is regulated through phosphorylation of NuMA by the centrosome-localized kinases PLK1 and CDK1 (Kiyomitsu and Cheeseman, 2012; Kotak et al., 2013; Seldin et al., 2013; Sana et al., 2018). Defects in bipolar spindle orientation have been observed when cortical dynein localization is perturbed via enhanced phosphorylation of NuMA or experimental depletion of LGN or Afadin (Kiyomitsu and Cheeseman, 2012; Carminati et al., 2016).

Cells with more than two centrosomes, termed centrosome amplification, are common in cancer. Extra centrosomes induce the formation of multipolar spindles and cell division can lead to the formation of multiple daughter cells that are nonviable (Kwon et al., 2008). However, cells with centrosome amplification often cluster their extra centrosomes into a functional bipolar spindle (Figure 1C), enabling a bipolar division and giving rise to two viable daughter cells (Kwon et al., 2008). This clustering activity is dependent on both passive and active mechanisms. Passive mechanisms, like contractility of the cortex-associated actin cytoskeleton, contribute broadly to centrosome positioning within the cell (Rhys et al., 2018). This helps to bring centrosomes within proximity of each other, where active mechanisms - those that allow centrosomes to engage and move with respect to each other, become relevant. Both motor-derived forces and crosslinking activity at the centrosomes have been shown to impact active clustering (Quintyne et al., 2005; Kwon et al., 2008; Barr and Gergely, 2008; Fielding et al., 2010; Leber et al., 2010; Ding et al., 2017). While cortical dynein-driven centrosome movement has been demonstrated to impact centrosome and spindle positioning in cells with 2 centrosomes, whether this activity may be relevant for either passive or active centrosome clustering in cells with centrosome amplification has not been investigated.

**Figure 1:**
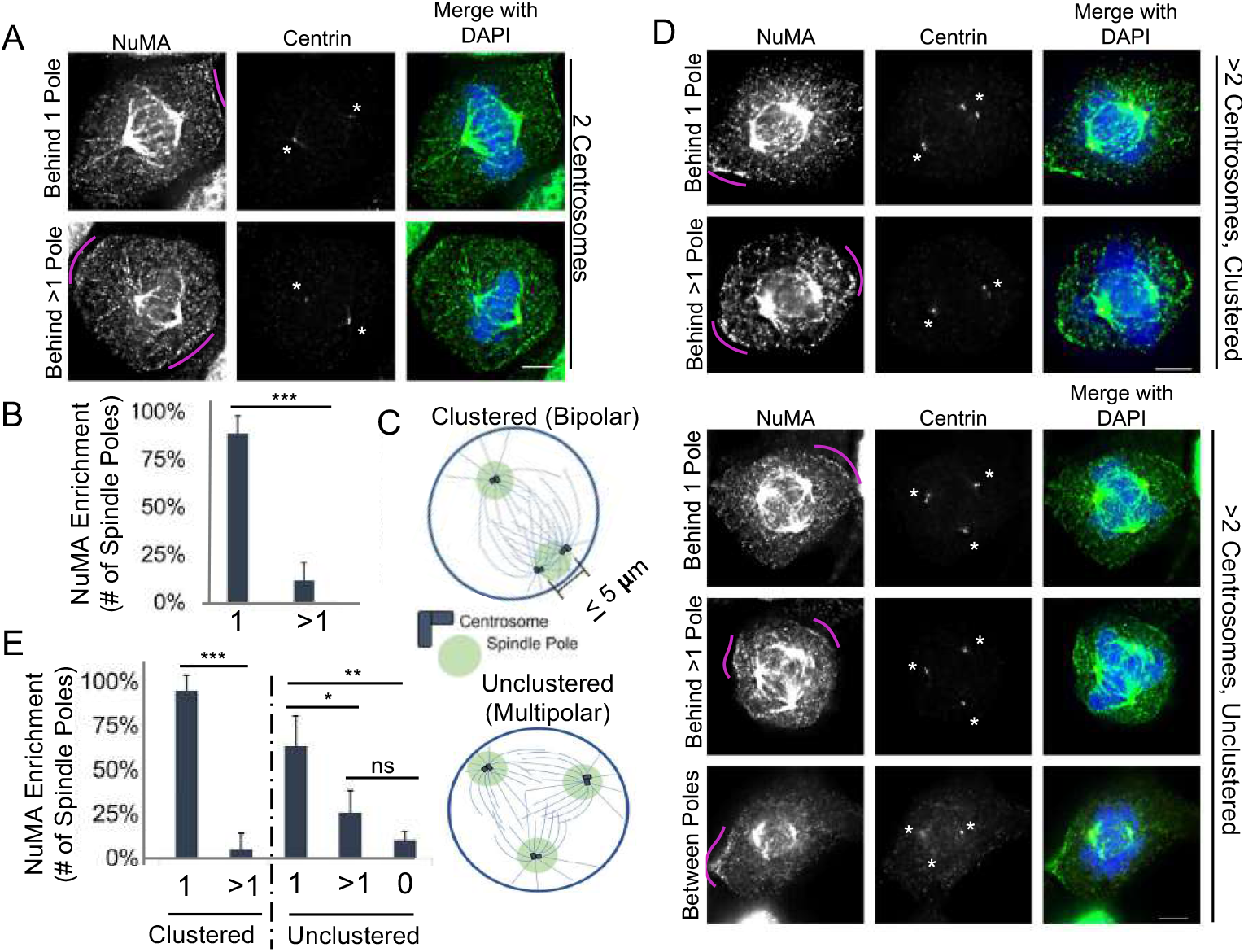
NuMA is primarily enriched in a region behind a single spindle pole, regardless of centrosome number. (A,B) Representative fixed-cell images and quantification of cortical NuMA enrichment in mitotic RPE cells with two centrosomes. NuMA was characterized as being behind 1 spindle pole or > 1 spindle pole. (C) Schematic of cells with three centrosomes that cluster and form a bipolar spindle (top) or are unclustered and form a multipolar spindle (bottom). Centrosome clustering is defined as a spindle pole with two or more centrosomes that are within 5 *μ*m apart. (D,E) Representative fixed-cell images and quantification of cortical NuMA enrichment in mitotic RPE cells following DCB treatment, which induces cytokinesis failure and results in cells with >2 centrosomes. NuMA was characterized as being behind 1 spindle pole, > 1 spindle pole, or between poles (behind 0 poles). All RPE cells are stained with antibodies specific for NuMA (green) and centrin-2 (red); DNA is detected with DAPI (blue). In (A) and (D), pink arcs highlight regions of increased cortical NuMA, white asterisks denote spindle poles, and scale bars are 5 *μ*m. In (B) and (E), error bars are standard deviation. In (E), cells were grouped based on spindle polarity and analysis performed on 50 cells per group for each of 3 biological replicates. Significance in (B) and (E) is determined by student’s t-test when comparing two conditions and a one-way ANOVA when comparing multiple conditions; \**p* < 0.05,\*\**p* < 0.01, \*\*\**p* < 0.001, ns indicates not significant.

We exploit a combination of molecular manipulation and computational cell modeling approaches to quantify the impact of cortical dynein-derived forces on centrosome clustering. We utilize image-based approaches to determine the localization of cortical NuMA/dynein in cells with centrosome amplification and assess how disruption of cortical dynein impacts centrosome clustering and mitotic progression. Informed by our experimental results, we define a computational model of mitotic spindle formation and function to explore the relationship between cortical dynein-derived forces and centrosome movement in cells with centrosome amplification. Our findings implicate cortical dynein in the directed centrosome movement necessary to efficiently cluster centrosomes in cells with centrosome amplification.

## Results

### Cortical Dynein Promotes Centrosome Clustering

In cells with two centrosomes, dynein and its co-factor NuMA are localized to the cell cortex in an asymmetric manner such that enrichment is primarily observed behind one spindle pole at a time (Kiyomitsu and Cheeseman, 2012). To assess whether similar localization is seen in cells with more than two centrosomes, we first induce centrosome amplification in Retinal Pigment Epithelial (RPE) cells through induction of cytokinesis failure (Supplemental Figure S2 C) by treating cells with dihydrocytochalasin B (DCB). This approach generates tetraploid cells with four centrosomes (Andreassen et al., 1996). We next performed immunofluorescent imaging to assess NuMA localization. NuMA functions to link dynein to the actin cytoskeleton and its localization mirrors that of dynein at the cell cortex (Kiyomitsu and Cheeseman, 2012; Seldin et al., 2013). Consistent with published work (Kiyomitsu and Cheeseman, 2012; Seldin et al., 2013), we find that cells with two centrosomes primarily exhibit NuMA enrichment behind a single pole (Figure 1 A,B). Similarly, we find that NuMA is also primarily enriched behind a single spindle pole in cells with centrosome amplification (Figure 1 D,E). This is true regardless of whether centrosomes are clustered together to form a bipolar spindle (where one or more spindle poles have two or more clustered centrosomes, Figure 1 C), or remain unclustered to form a multipolar spindle.

Work from our group and others has implicated cortical dynein as a dominant driver of centrosome movement during mitosis in cells with two centrosomes (di Pietro et al., 2016; Mercadante et al., 2021). To test whether cortical dynein-driven centrosome movement is involved in centrosome clustering in cells with centrosome amplification, we perturbed dynein localization or function in cells with experimentally induced centrosome amplification. Two complementary approaches were used to induce centrosome amplification: overexpression of PLK4 (indPLK4) to induce the biogenesis of extra centrosomes (Godinho et al., 2009) or induction of cytokinesis failure, as described above (Supplemental Figure S2 A,C). Next, cortical dynein motor activity was disrupted using the small molecule inhibitor dynarrestin (Hoing et al., 2018; Mercadante et al., 2021), or dynein localization was perturbed via siRNA-mediated depletion of Afadin or LGN (B,D). Individual cells were analyzed for centrosome number, centrosome location, and spindle structure, where clustering was assessed pre-anaphase and bipolar spindles were considered as those in which one or both spindle poles contained two or more centrosomes located within 5 *μ*m of each other (Figure 2 A,B). Cells with centrosome amplification and functional cortical dynein are efficient at clustering centrosomes, such that ~50% of cells with >2 centrosomes are able to form clustered bipolar spindles (Figure 2 D,E). In contrast, inhibition of dynein (dynarrestin) or dynein delocalization from the cortex (Afadin or LGN depletion) decreased centrosome clustering such that only ~20% of cells with centrosome amplification exhibit clustered bipolar spindles (Figure 2 D,E).

**Figure 2:**
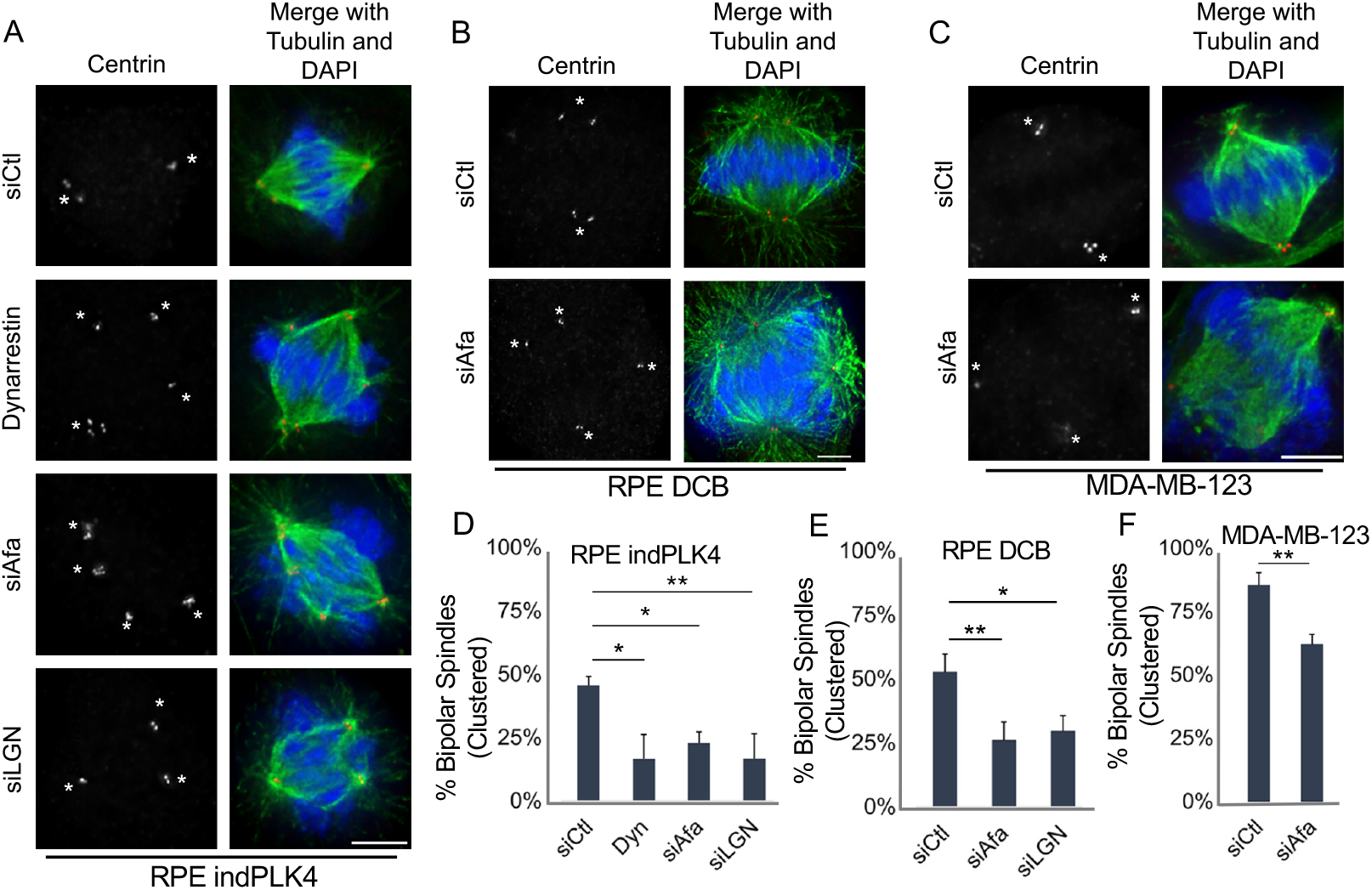
Centrosome clustering is compromised in the absence of cortical dynein activity. Representative fixed-cell images of cell lines with extra centrosomes, either through experimental manipulation inducing overexpression of PLK4 (A: RPE indPLK4) or treatment with DCB to induce cytokinesis failure (B: RPE DCB), or preexisting centrosome amplification (C: MDA-MB-31). Cells were treated with the dynein inhibitor dynarrestin, or treated non-targeting (siCtl) or LGN or Afadin-targeting siRNA (siLGN or siAfadin) to disrupt dynein localization at the cortex (see Supplemental Figure S2 for experimental setup and confirmation of Afadin and LGN depletion). Antibodies specific for alpha tubulin (green), centrin-2 (red), and DAPI (blue) to detect DNA were utilized to assess mitotic spindle structure. The white asterisks denote spindle poles, and scale bars are 5 *μ*m. Quantification of the percent of mitotic cells (pre-anaphase) exhibiting clustered bipolar spindles are given in D (RPE indPLK4), E (RPE DCB), and F (MDA-MB-31). All analysis was performed on at least 50 cells per condition from 3 biological replicates and rrror bars are standard deviation. Significance was determined via a student’s *t*-test when comparing two conditions and a one-way ANOVA with Dunnett’s test for multiple comparisons when comparing multiple conditions to one control; \**p* < 0.05,\*\**p* < 0.01.

To explore whether dynein activity similarly impacts centrosome clustering in cells with pre-existing centrosome amplification, we perturbed cortical dynein localization in MDA-MB-231 cells (Supplemental Figure S2 E,G), a well characterized breast cancer cell line with centrosome amplification (Mittal et al., 2017). Centrosome positioning and spindle structure was assessed as described above. Nearly 45% of mitotic MDA-MB-231 cells have centrosome amplification (Supplemental Figure S2 F). Of the mitotic cells with greater than two centrosomes, ~80% exhibit centrosome clustering and bipolar spindle formation (Figure 2 C,F). Consistent with our results in cells with induced centrosome amplification, MDA-MB-231 cells exhibit a reduction in centrosome clustering when cortical dynein localization is perturbed, such that only ~60% of pre-anaphase cells form a clustered bipolar spindle following Afadin depletion (Figure 2 F).

### Cells Complete Multipolar Divisions in the Absence of Cortical Dynein

Centrosome clustering is a dynamic process that can take an hour or longer to achieve (Kwon et al., 2008; Navarro-Serer et al., 2019) and analysis of a single time point in mitosis cannot distinguish between a delay versus a deficit in centrosome clustering. To further assess the observed increase in multipolar spindles seen following cortical dynein disruption, cells were additionally treated with the protease inhibitor MG132 prior to assessment of centrosome positioning and spindle structure. MG132 prevents anaphase onset, which provides cells additional time in which to cluster centrosomes. Consistent with previous findings (Rhys et al., 2018), cells with centrosome amplification progress from ~50% to 60% and then 80% bipolar spindles following 30 or 60 minutes of MG132-induced mitotic arrest, respectively (siCtl in Figure 3 A,B). In contrast, the percent of mitotic cells with extra centrosomes and perturbed dynein activity that are able to form a bipolar spindle does not increase with prolonged mitotic duration. Instead, cells with centrosome amplification remain primarily multipolar, even after 60 minutes of MG132 treatment (Figure 3 A,B).

**Figure 3:**
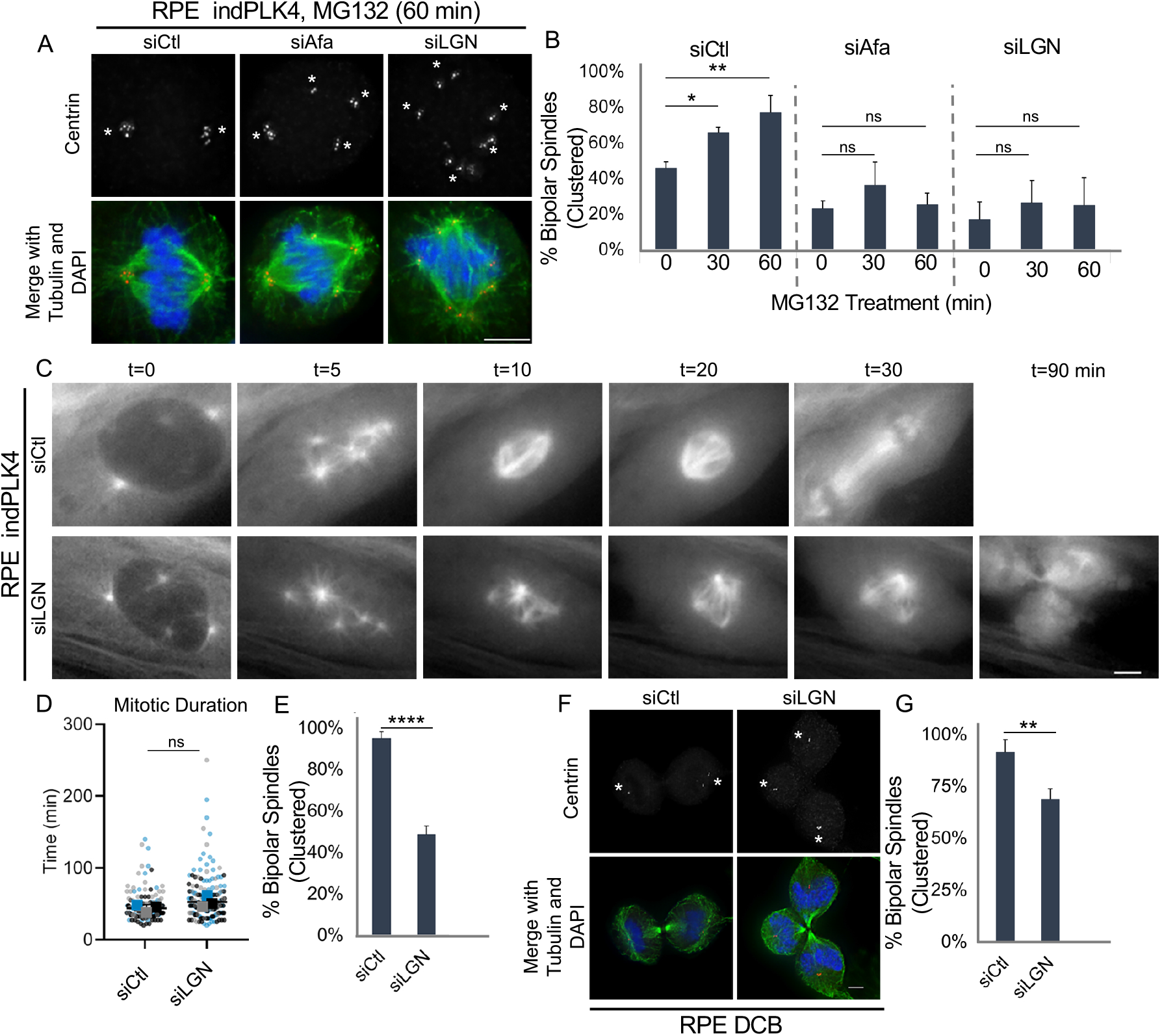
Cortical dynein activity promotes centrosome clustering and bipolar cell division. (A,B) Representative fixed-cell images and quantification of the frequency of spindle bipolarity in cells induced to have extra centrosomes through overexpression of PLK4 (RPE indPLK4). Cells were treated with non-targeting (siCtl) or Afadin or LGN-targeting (siAfa or siLGN) siRNA to disrupt cortical dynein localization, subsequently treated as indicated with MG132 to prevent anaphase onset. Antibodies specific for centrin-2 (red), tubulin (green), and DAPI (blue) to detect DNA were utilized to assess spindle structure. (C) Still frames from live cell imaging of RPE indPLK4 cells expressing EGFP-tubulin are shown for both control (siCtl) and LGN (siLGN) depleted cells at indicated timepoints. (D) Mitotic progression of RPE indPLK4 cells were timed from nuclear envelope breakdown (characterized by loss of EGFP exclusion from the nucleus) until anaphase B (indicated by rapid elongation of the spindle) and (E) the frequency of mitotic cells progressing through a bipolar division was quantified. (F,G) Representative images and quantification of cells progressing through a bipolar division (determined at anaphase/telophase) in cells with extra centrosomes acquired through DCB-induced cytokinesis failure. The control and LGN depleted RPE DCB cells were stained with antibodies specific for centrin-2 (red) and tubulin (green), and assessed for evidence of bipolar vs multipolar divisions. Scale bars in (A,C,F) are 5 *μ*m. All quantifications were performed on at least 50 cells per condition from each of 3 biological replicates and error bars are standard deviation. Significance was determined by student’s *t*-test when comparing two conditions and a one-way ANOVA with Dunnett’s test for multiple comparisons when comparing multiple conditions to one control; \**p* < 0.05,\*\**p* < 0.01, \*\*\*\**p* < 0.0001, ns indicates not significant.

To assess the temporal relationship between spindle dynamics and mitotic fate of cells following loss of cortical dynein activity, we performed timelapse imaging of cells stably expressing fluorescently-tagged tubulin (EGFP-tubulin) (Figure 3C) (Mercadante et al., 2019) and induced cells to have extra centrosomes through induction of PLK4 expression, as described above. Phase contrast and fluorescence images were captured every 2.5 minutes throughout mitosis and used to assess spindle structure, mitotic timing, and the number of progeny resulting from each division. We find that individual mitotic cells with centrosome amplification cluster extra centrosomes and progress through mitosis in ~45 min. The vast majority (95%) of these cells complete a bipolar division (Figure 3 D,E). Disruption of cortical dynein activity does not alter mitotic timing (Figure 3 D). However, consistent with centrosome clustering defects described above, 50% of cells with centrosome amplification that have disrupted cortical dynein activity exit mitosis with a multipolar division (Figure 3 D,E). Fixed-cell analysis reveals similar results in anaphase and telophase cells with centrosome amplification induced via DCB (Figure 3 F,G). Together, these results implicate cortical dynein as necessary for centrosome clustering and bipolar division in cells with centrosome amplification.

### Centrosome Clustering is Sensitive to Dynein Activity at the Cortex

To further manipulate cortical dynein and explore the extent to which it impacts centrosome clustering in cells with centrosome amplification, we developed a force balance model that tracks centrosome movement and spindle formation, taking into account major motor-derived forces during mitosis. This model was informed by our previous work (Mercadante et al., 2021) and modified to account for the presence of >2 centrosomes. To simulate a cell that has rounded in mitosis, the cell boundary is defined as a rigid circle with a diameter of 30 *μ*m. Dynamic microtubules are short at mitotic entry and elongate through mitotic progression (Figure 4A). Motor proteins Eg5, HSET, and dynein push or pull on the microtubule (and hence exert force on the centrosome the microtubule is attached to). These motor-dependent forces, along with forces associated with microtubules pushing on the cell cortex, are evaluated at 0.5 second intervals to determine centrosome movement in the mitotic cell (Supplemental Figure S1 A). Motor-dependent force generation is dependent on distances between model entities and modulated by altering the stochastic binding probability of a motor; increased binding probability equates to increased motor activity (Supplemental Figure S1 B). Additional model details are given in the Materials and Methods Section.

**Figure 4:**
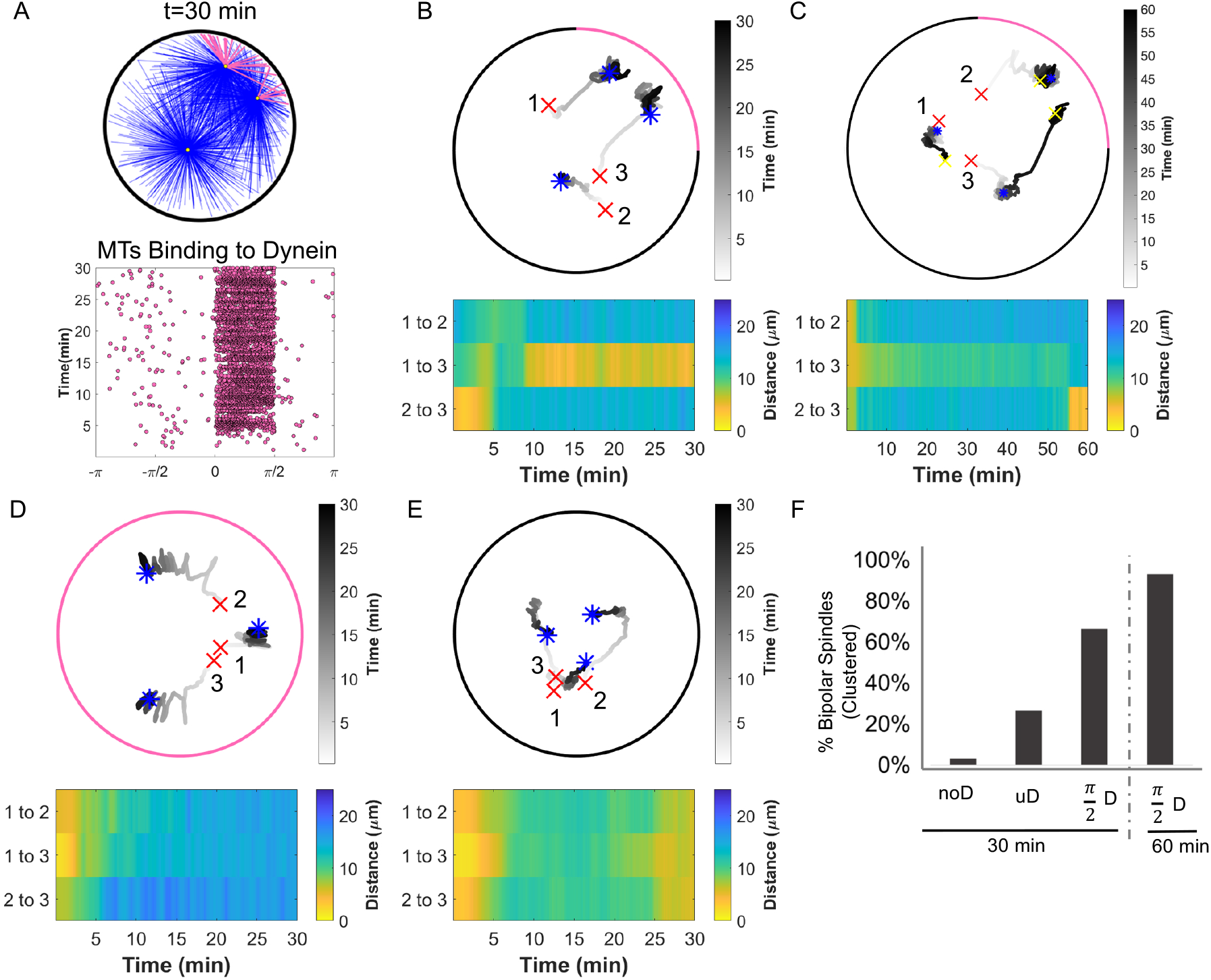
Restricted cortical dynein localization drives centrosome clustering. (A) Simulation results where cortical dynein is enriched in the region from 0 to *π*/2. (A, Top) Simulation at t = 30 min showing centrosomes in yellow, microtubules bound to cortical dynein in pink, and all others microtubules in blue. (A, Bottom) Plot of microtubules binding to cortical dynein on the boundary of the cell (–*π* to *π*) from the simulation shown above. Each dot indicates an individual microtubule binding to dynein. (B-E) Top: Trace of centrosome movement over time through the duration of a simulation where a red ‘x’ indicates initial centrosome position, a blue ‘*’ indicates centrosome position at 30 minutes, and a yellow ‘x’ (C only) indicates centrosome position at 60 minutes. Numbers mark individual centrosomes, grayscale indicates time, and pink on the cell boundary indicates the region of high dynein activity (where *P_d_cor__* = 0.5; elsewhere *P_d_cor__* = 0.01). Bottom: Heat map representing the pairwise distances between all centrosome pairs indicated in the corresponding traces in the top panel. (F) The percent of simulations that achieve bipolar spindles (centrosome clustering) when cortical dynein is absent (noD), distributed uniformly across the cell boundary (uD), or enriched in the region from 0 to *π*/2 ((*π*/2)D); duration of simulations are *t*=30 or 60 minutes as indicated. Data in (F) represent 30 simulations for conditions shown in (B,D,E) and 15 simulations for the condition shown in (C).

Our immunofluorescence analysis showed that regardless of centrosome number, dynein primarily localized in a region behind a single spindle pole (Figure 1 E,F). To assess the functional relevance of the NuMA/dynein distribution on centrosome clustering, we manipulated the distribution of cortical dynein activity in our model. The NuMA/dynein distribution was mimicked by permitting microtubules on centrosomes to bind to cortical dynein with probability *P_d_cor__* = 0.5 on one fixed quadrant (angular region of *π*/2), while having a much smaller binding probability, *P_d_cor__* = 0.01, everywhere else (Figure 4 A). We then assessed the movement of three centrosomes over time and quantified the pairwise distances between all centrosome pairs. In Figure 4 B, centrosomes numbered 2 and 3 initially start close together, but move apart within a few minutes whereas centrosomes 1 and 3 start further apart and cluster together around *t* = 10 min and remain clustered up until *t* = 30 min. We define this as sustained clustering-centrosomes that have a pairwise distance of ≤5 *μ*m that is maintained for the duration of the simulation after initial centrosome separation (after *t* = 5 min). This simulation is also classified as having formed a bipolar spindle where the sustained clustering of centrosomes 1 and 3 is one pole and the other pole is centrosome 2. In contrast, cases where centrosomes stay 7 to 20 *μ*m apart after *t* = 5 min and do not cluster are shown in Figure 4 D and E, corresponding to the case of uniform dynein (*P_d_cor__* = 0.5 everywhere on the cortex) and no dynein (*P_d_cor__* = 0.01 everywhere on the cortex), respectively.

Quantification of cortical dynein-derived forces when dynein has fixed enrichment in a region reveals a correlation between centrosome clustering and cortical dynein. Peaks in dynein-derived forces from pulling on microtubule tips at the cortex coincides with significant decreases in the pairwise distances between centrosomes to which those microtubules are anchored for the case of dynein in an angular region of *π*/2 (Supplemental Figure S5 A,E). Two centrosomes generally exhibit sustained clustering and form a pole closer to the region of dynein enrichment whereas the other centrosome becomes the other pole and is ~17 *μ*m from the enriched region and not interacting with cortical dynein (Supplemental Figures S5 C,D and S6 A). With dynein uniformly distributed on the cortex, we observe cortical dynein acting on microtubules associated with all centrosomes; all centrosomes end up at a distance of ~5 *μ*m from the cortex, but are not able to cluster (Supplemental Figures S5 B,C and S6 B). In the case of no dynein, centrosomes do not cluster and pairwise distances between centrosomes stabilizes at ‘8 *μ*m apart (Supplemental Figures S5 D,E and and S6 C).

Due to the inherent stochasticity of motor and microtubule dynamics, we ran 30 simulations (unless otherwise stated) to quantify the frequency of sustained clustering and bipolar spindle formation. We find that in simulations with three centrosomes and cortical dynein primarily in a single region or quadrant, ~65% of cells formed bipolar spindles within 30 minutes (Figure 4, A,B,F). Bipolar spindle formation is greatly reduced to ~25% and ~5% in the case of uniformly distributed dynein and no dynein, respectively (Figure 4 F). The reduction in bipolar spindle formation in the case of no dynein is consistent with our biological results in RPE cells with centrosome amplification and perturbed dynein localization (Figure 2 D-F). Given that our biological experiments indicate that the frequency of centrosome clustering is sensitive to mitotic duration (when cortical dynein is not perturbed), we next sought to determine if centrosome clustering in our model is similarly sensitive to simulation duration. Consistent with our biological results, extension of the simulation from 30 to 60 minutes increases the frequency of centrosome clustering from ~65% to ~90% (Figure 3 A,B; Figure 4 C,F).

### Oscillations in Cortical Dynein Enrichment Impact Centrosome Movement

In cells with two centrosomes, cortical dynein localization is primarily behind one spindle pole at a time and oscillates from behind one spindle pole to the other every few minutes (Kiyomitsu and Cheeseman, 2012). To determine if dynamic cortical dynein localization influences centrosome clustering in cells with centrosome amplification, we mimicked the observed biological oscillations by moving the *π*/2 angular region of enrichment to the opposite side of the cortex every *T* minutes (*T* = 5, 10, and 15 in Figure 5A-C and *T* = 1.67, 2.33 in Supplemental Figure S7 A,B). In contrast to static dynein enrichment in an angular region of *π*/2, we do not observe sustained clustering for periods of 20 or 25 minutes (comparing Figure 4B with Figure 5A-C). Instead, centrosomes with microtubules bound to cortical dynein moved toward the region of dynein activity, effectively dragging similarly engaged centrosomes towards each other and when dynein activity oscillates to the opposing quadrant, previously engaged centrosomes do not sustain their close proximity to each other (Figure 5A-C; Supplemental Figure S7 B).

**Figure 5:**
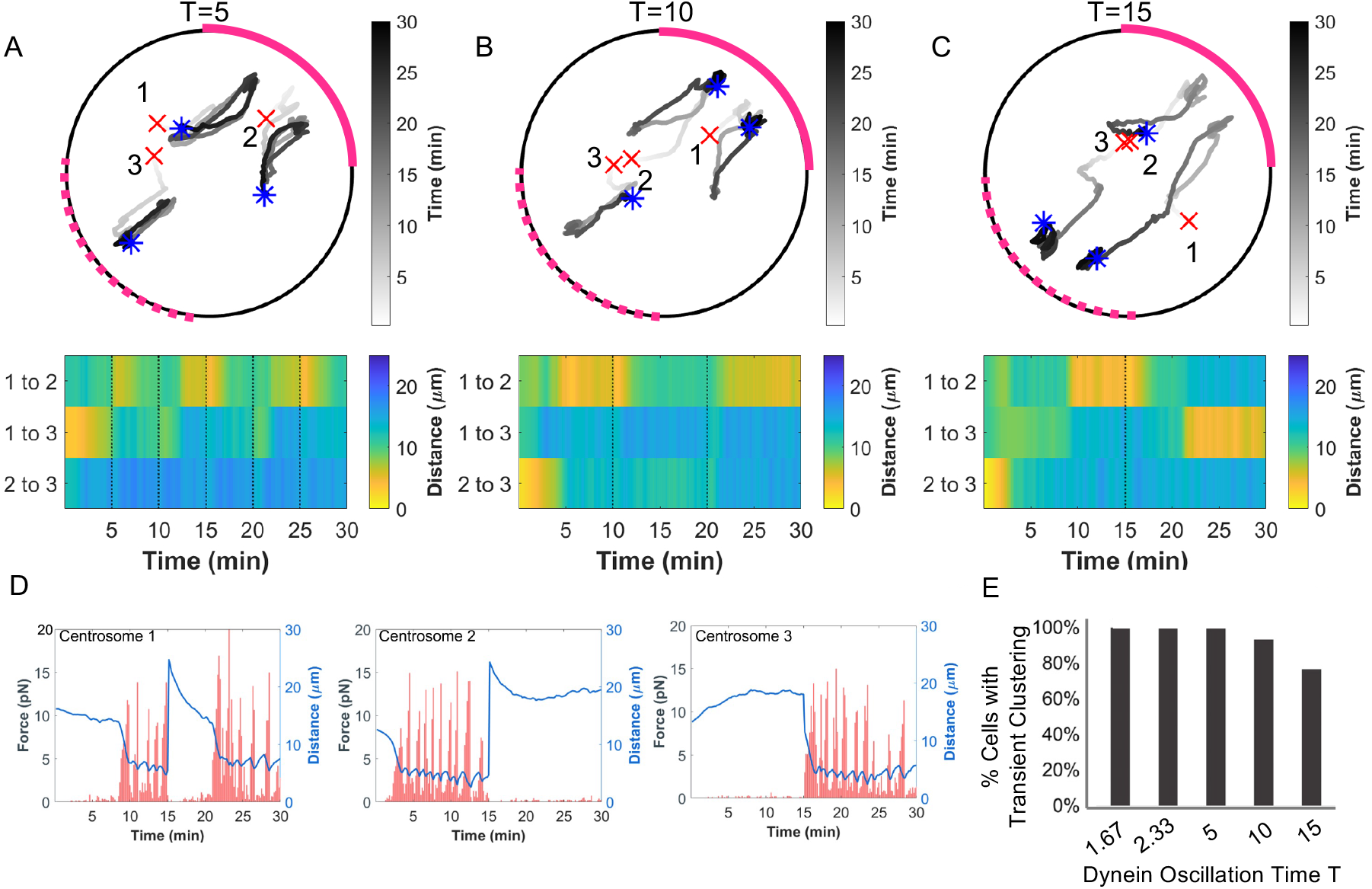
Oscillatory redistribution of cortical dynein activity enhances centrosome clustering. (A,B,C) Top: Traces of centrosome movement over time from a simulation with dynein enriched regions (with *P_d_cor__* = 0.5) oscillating between the upper-right (0 to *π*/2, solid pink arc) and lower-left (–*π* to –*π*/2, dashed pink arc) quadrants with a period of T = 5 min (A), 10 min (B) or 15 min (C). Initial centrosome position indicated by a red ‘x’ and final centrosome position indicated by a blue ‘*’. Grayscale indicates time. Bottom: Heat map representing the pairwise distances between all centrosomes from the corresponding simulation above. Black dotted lines indicate a timepoint when cortical dynein localization is redistributed. (D) Plots depicting the magnitude of cortical dynein-derived forces over time (red trace) and the distance between each centrosome and the mid-point of dynein localization (blue trace) from the simulation shown in (C), where cortical dynein is redistributed at 15 min. (E) Quantification of transient centrosome clustering at different periods of cortical dynein oscillations; 30 simulations for each condition shown in (B,D,E) and Supplemental Figure S7 A,B.

We do observe transient clustering where centrosomes are ≤5 *μ*m apart for several minutes and only move apart after the region of dynein localization oscillates to the other side of the cortex. For example, in Figure 5 C, in the first 15 minute period where dynein is enriched in the upper right quadrant of the cortex, centrosomes 1 and 2 exhibit clustering and cortical dynein-induced forces on these centrosomes is largest for *t* = 10 – 15 minutes (Figure 5 D). At *T* = 15 minutes, dynein is then enriched in the lower left quadrant and we observe the pairwise distances between centrosomes 1 and 2 increasing whereas the pairwise distances of centrosomes 1 and 3 decrease and they cluster (with increased cortical dynein-induced forces acting on these centrosomes after t = 20 minutes, Figure 5 D). To determine whether changes in pairwise centrosome distance is a consequence of loss of cortical dynein behind the clustered pair or instead results from dynein-dependent pulling from the opposing side, we performed simulations in which microtubules bind to asymmetric cortical dynein until *t* = 15 minutes and then dynein is removed for the duration of the simulation (*P_d_cor__* = 0.01 everywhere for *t* ≥ 15 minutes). Assessment of dynein-derived forces and corresponding centrosome movement reveals that centrosomes cluster toward the region of enriched dynein, but begin separating after dynein is removed at *t* = 15 minutes (Supplemental Figure S7, C and D), indicating that loss of clustering is a passive event.

To quantify the propensity for a bipolar spindle to form, we recorded the number of simulations where we observe transient centrosome clustering at least one time, sustained for at least one and a half minutes. The frequency of transient centrosome clustering in simulations with cortical dynein oscillations is >80%, regardless of the period of dynein oscillations (Figure 5E). This propensity to form a bipolar spindle is much higher than ~60% of simulations achieving centrosome clustering when cortical dynein activity was stably enriched in one quadrant of the cortex (comparing to Figure 4 F). Together, these results suggest that centrosome movement and clustering is driven by cortical dynein activity.

## Discussion

In this work, we combine biological experimentation with computational modeling to inform novel dynein-dependent mechanisms driving centrosome clustering in cells with extra centrosomes. Our simulations indicate that cortical dynein-dependent forces are responsible for actively directing movement of centrosomes toward the cell cortex (Figure 4; Supplemental Figure S5). When dynein localization along the cell cortex is primarily in a region behind one of the poles of the mitotic spindle (Figure 1), dynein-dependent forces serve to bring individual centrosomes within close proximity of each other, near the region of cortical dynein activity (Figures 4 and 5; Supplemental Figures S3 and S7). We propose that this directed movement of centrosomes toward a common region on the cell cortex is indicative of active centrosome clustering. The centrosome(s) not near the region of cortical dynein activity do not cluster and instead are held at a stable distance from clustered centrosomes by other microtubule derived forces (i.e. binding to dynein at spindle poles, Eg5, and/or HSET, and pushing on the cell cortex; Supplemental Figure S6), resulting in a clustered bipolar spindle structure (Figures 4 and 5; Supplemental Figures S3 and S7). Centrosome clustering in this model is dependent on cortical dynein activity and is sensitive to the distribution of dynein on the cell cortex, as either removing cortical dynein activity from simulations or defining dynein to be uniformly distributed on the cortex precludes centrosome clustering (Figure 4; Supplemental Figure S4). Our biological data support this model and demonstrate that cortical dynein contributes to centrosome clustering in cells with centrosome amplification, such that loss of cortical dynein results in primarily unclustered spindles and multipolar divisions (Figures 2 and 3).

Our modeling framework provides a way to exploit temporal and spatial regulation of cortical dynein activity in ways not easily achieved in biological systems. For example, regions of primary dynein localization varied in size in our cell lines (Figure 1 E,F), and we can explore how the size of this region may impact the propensity of centrosomes to cluster. Simulations assessing progressively smaller angular regions of high cortical dynein activity from *π* to *π*/8 revealed that the frequency of clustering is maximized at *π*/2 and decreases as the region of dynein activity is increased above or decreased below *π*/2 (Supplemental Figure S4). Centrosome clustering and bipolar spindle formation is found to be sensitive to the size of the fixed region where cortical dynein is enriched in cells with centrosome amplification. Additionally, cells with induced or naturally occurring centrosome amplification exhibit a range of centrosome numbers and may have different regions of cortical dynein enrichment. In our model, we can fix dynein enrichment to an angular region of *π*/2 and explore dynein-induced centrosome movement in simulated cells with 2, 3, or 4 centrosomes. Cortex-directed centrosome movement did correspond with peaks in dynein activity (Supplemental Figure S5) for simulations with two and more than two centrosomes (Supplemental Figure S8; Supplemental Figure S5) (Mercadante et al., 2021). Our base model of mitotic spindle formation with supernumerary centrosomes reproduces critical features of passive centrosome clustering and is robust to centrosome number-revealing similar frequency and dynamics of spindle pole clustering with both three and four centrosomes with fixed dynein enrichment as well as perturbations to dynein localization (Figure 4 and Supplemental Figure S3). These results show that the propensity to form a bipolar spindle is sensitive to the region of dynein localization whereas formation of a bipolar spindle with a given region of dynein enrichment is not sensitive to centrosome number.

Previous studies of mitotic cell division utilizing different modeling approaches have been valuable in understanding and informing force-derived centrosome clustering mechanisms in cells with centrosome amplification (Chatterjee et al., 2020; Goupil et al., 2020; Miles et al., 2022). In particular, these models have provided insight into chromosome-dependent centrosome clustering mechanisms which implicated kinetochore microtubule-derived torque (Miles et al., 2022) and have highlighted that there must exist a delicate balance between attraction forces for efficient centrosome clustering to occur, including centrosome-cortex forces (Chatterjee et al., 2020). Our results further expand on this latter observation by identifying that the centrosome-cortex force must correspond to a region on the cell cortex, either fixed or dynamically changing, for efficient clustering to occur via dynein motor activation (Figure 4).

Chromosomes and chromosome-derived forces have been implicated in centrosome clustering in two distinct ways. First, chromosomes form a physical barrier that segments the cell, thereby restricting centrosome movement (Goupil et al., 2020). Second, chromosomes form stable interactions, via kinetochores, with bundles of microtubules that are in turn anchored at the centrosomes (DeLuca et al., 2006). The bioriented configuration, and associated forces, of paired kinetochores enforce a bipolar geometry where centrosomes are positioned along the spindle axis (Leber et al., 2010; Tanaka, 2010; Chatterjee et al., 2020; Miles et al., 2022). Similarly, cell shape and actin-dependent cortical contractility impact centrosome clustering by restricting the space within which centrosomes can move (Kwon et al., 2008; Rhys et al., 2018). Dynein is functionally linked to the actin cell cortex through the proteins NuMA/LGN/Afadin (di Pietro et al., 2016), suggesting that cortical dynein and actin-dependent contractility act in concert to drive passive clustering by bringing centrosomes into proximity of each other. Once centrosomes are within ~8 *μ*m of each other, activity by static crosslinkers and motor proteins (i.e. HSET) at spindle poles engage centrosome-associated microtubules and drive robust and sustained centrosome clustering (Quintyne et al., 2005; Kwon et al.,2008; Rhys et al., 2018). Our simulations lack both chromosome-derived forces and crosslinking activity, explaining why when cortical dynein activity is turned off or redistributed on the cell cortex, previously clustered centrosomes begin to move apart (Figure 5F).

Our results indicate cortical localization of dynein alone is not sufficient for centrosome clustering and instead specific enrichment of dynein behind a single spindle pole is critical to drive clustering.

Since cortical dynein localization is negatively regulated by centrosome-localized kinases (Kiyomitsu and Cheeseman, 2012), this suggests an iterative process to achieve and sustain centrosome clustering where dynein pulls centrosomes towards a common point on the cell cortex and then is itself inhibited by the centrosomes clustered nearby. We propose that regional enrichment of cortical dynein can both drive and be driven by centrosome clustering. Additional mechanisms such as kinase activity, non-motor activity, and tension from kinetochore-microtubule interactions would then allow for sustained clustering when regions of dynein localization are dynamic.

The presence of centrosome amplification is a hallmark of cancer and is associated with drug resistance, tumor progression, and poor patient prognosis (D’Assoro et al., 2002; Fukasawa, 2005; Mittal et al., 2020). Due to the requirement of cancer cells with centrosome amplification to cluster their centrosomes to remain proliferative, prevention of centrosome clustering in cancer cells is believed to be a promising therapeutic approach (Kwon et al., 2008; Leber et al., 2010; Godinho and Pellman, 2014; Sabat-Pospiech et al., 2019). The motor protein HSET, along with proteins involved in the formation and maintenance of cell-cell junctions, cortical contractility, and kinetochore-microtubule interactions have been implicated as potential targets to limit centrosome clustering in cancer cells (Kwon et al., 2008; Hebert et al., 2012; Kwon et al., 2015; Rhys et al., 2018). Dynein had previously been implicated in centrosome clustering, although this role was believed to be associated with its function at spindle poles (Quintyne et al., 2005). Our data now indicate that both cortical dynein activity and its dynamic and asymmetric localization behind spindle poles are critical for efficient centrosome clustering. Through simulations and molecular manipulations in cells with experimentally induced centrosome amplification or cancer cells with preexisting centrosome amplification, we show that perturbation of dynein’s ATP-driven motor activity or kinase-sensitive cortical localization impacts centrosome clustering. These results suggest that inhibition of either feature of dynein (activity or dynamic localization) may be of therapeutic interest in cancers with a high frequency of centrosome amplification.

## Materials and Methods

### Modeling

In this work, we optimize and expand upon our previously described two-dimensional model with dynamic microtubules and stochastic motor-dependent force generation to capture the centrosome movement in cells with centrosome amplification (Mercadante et al., 2021). We allow microtubule-motor protein interactions with Eg5 and HSET on antiparallel microtubules, capturing the dominant roles of these proteins in mitosis. Dynein is localized at the cell cortex and spindle poles to account for its functions in pole focusing and spindle dynamics, respectively (Supplemental Figure S1A) (Ferenz et al., 2009, 2010; van Heesbeen et al., 2014; Loncar et al., 2020; Mann and Wadsworth, 2019; Sharp et al., 2000; She and Yang, 2017; Vaisberg et al., 1993). Kinesin-5, Eg5 in mammalian cells, is a homotetrameric motor protein with two motor domains on either side of an elongated stalk (Kashina et al., 2009). Each of the motor domains binds to a microtubule and walks towards the plus-end (Bodrug et al., 2020). When the microtubules are antiparallel, as they are in the interpolar region of the spindle, this movement causes microtubule sliding in opposite directions and drives centrosome separation and early spindle formation (Kapoor et al., 2000; Mayer et al., 1999). Kinesin-14, HSET in mammalian cells, is a minus-end directed dimeric motor protein with two motor heads on one end of the molecule and non-motor microtubule binding domains on the other (Braun et al., 2009; Fink et al., 2009). At the interpolar region of the spindle, HSET facilitates antiparallel microtubule-microtubule sliding to help maintain mitotic spindle length (Fink et al., 2009). HSET movement opposes that of Eg5, resulting in an inward force between spindle poles (Mountain et al., 1999; Sharp et al., 2000). Dynein is the major minus-end directed motor protein in mammalian cells. While dynein at the cell cortex is essential for spindle positioning and orientation, it has additional essential roles within the spindle that are required for the maintenance of spindle structure. Dynein activity in the interpolar region of the spindle, where interpolar microtubules overlap, counteracts that of Eg5, with one of dynein’s two motor heads walking along each microtubule. This movement pulls spindle poles together, antagonizing centrosome separation and bipolar spindle formation (Ferenz et al., 2009; Raaijmakers et al., 2013; Raaijmakers and Medema, 2014). Additionally, dynein motor activity on parallel microtubules is critical for maintaining microtubule minus-end focusing at spindle poles, where loss of dynein results in splayed poles and barrel-like spindles (Echeverri et al., 1996; Goshima et al., 2005).

### Dynamic Microtubules

Microtubules emanate from a centrosome and are initialized with uniformly distributed random lengths *ℓ* and angles *α* (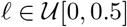 *μ*m, and angle 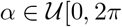). At each time step, new microtubules are randomly nucleated a rate *MT_nuc_*. Microtubules are modeled as semi-rigid filaments with a constant bending rigidity *κ*. Microtubule plus-ends are dynamic and have constant growth velocity *ν_g_* and shrinking velocity that is *ν_b_* for microtubules bound to dynein and *ν_s_* for all other shrinking microtubules. Microtubule dynamic instability for each microtubule *i* is defined by a constant rescue frequency (*k*_1_) and a length-dependent catastrophe frequency (*k*_2_*i*__). We use a stochastic Monte-Carlo method to determine microtubule dynamic instability; a random number 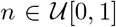 is generated and if *n* ≤ 1 –*e*^−*k*_1_*dt*^ or *n* ≤ 1 –*e*^–*k*_2_*i*__*dt*^, the microtubule will undergo rescue or catastrophe, respectively (Supplemental Figure S1B). Shrinking microtubules that do not undergo rescue will depolymerize completely and no longer be considered in the system when *ℓ_i_* ≤ *ν_g_*dt**. Throughout the simulation, the microtubule length *ℓ* and angle *α* are updated based on the state of the microtubule (Supplemental Figure S1B).

### Initialization and Algorithm

Centrosomes are initialized in random positions within 7.5 *μ*m from the cell center, to simulate positioning following nuclear envelope breakdown at the start of mitosis. At each time step, new microtubules are nucleated for each centrosome at a rate *MT_nuc_*. Microtubule dynamics and microtubule binding to motor proteins are determined and updated by a set of stochastic rules based on distances and probabilities (Supplemental Figure S1B). The force on each microtubule is calculated and summed to determine the total force on centrosome c. The position of centrosome c is updated based on Equation 4. The length and angle of each microtubule is updated based on its state and this process is repeated until *t* = 30 minutes. Parameters are highlighted in Supplemental Table S1.

### Stochastic Microtubule-Motor Interactions

Microtubule dynamic instability results in changes in microtubule length and positioning at every time step. As such, microtubule interactions with motor proteins are transient, with motor interactions depending on proximity to a microtubule. In addition to proximity, we consider each motor protein population (Eg5, HSET, cortical dynein, and dynein at spindle poles) to have a distinct binding probability (Table S1). If a distance argument and probability are satisfied, the microtubule will bind to the motor protein. For example, a microtubule tip within a distance 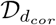 from the cell boundary will then bind to cortical dynein with probability *P_d_cor__* (implemented by choosing 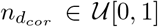 and allowing binding to occur if *n_d_cor__* < *P_d_cor__*). Otherwise, the microtubule will continue to grow and push against the cell boundary with a length-dependent force:

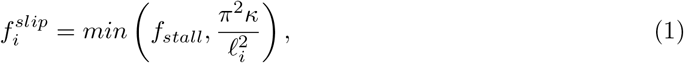

where *f_stall_* is the stall force of the microtubule and *κ* is the bending rigidity. Interpolar microtubules *i,j* nucleated from centrosomes *c, k*, that are within a distance 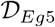 or 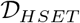 will have a probability of binding to Eg5 (*P_E_*) and/or HSET (*P_H_*) and generating force. Using a Monte Carlo Method, if a random number *n*_*Eg*5_, *n_HSET_* is less than *P_E_, P_H_*, binding of Eg5 and/or HSET occurs, respectively. We allow each microtubule from centrosome *c* to have Eg5 and/or HSET binding on up to two microtubules from centrosome *k*. Microtubules also interact with spindle pole dynein near opposing centrosomes when a microtubule gets within a distance 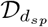 and satisfies the probability *P_d_sp__* (Supplemental Figure S1B).

### Motor-Dependent Force Generation and Centrosome Movement

We consider motor-dependent forces to be stochastic, where force by motor *m* is generated if both a distance argument between the two interacting structures (microtubule-microtubule, microtubule cortex, or microtubule-spindle pole) and a motor-specific binding probability *P_m_* are satisfied (Supplemental Figure S1B). Individual motor forces on the *i^th^* microtubule, 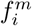, are calculated using a standard force-velocity relationship (Svoboda and Block, 1994):

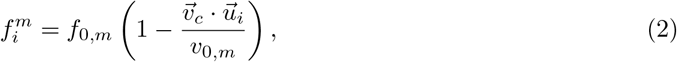

where *f*_0,*m*_ is the stall force of motor *m, ν*_0,*m*_ is the walking velocity of motor *m*, 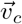 is the velocity of centrosome *c* that microtubule *i* emanates from, and 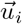 is the unit vector in the direction of microtubule *i*. The total force by all motors *m* bound to the *N_c,m_* microtubules nucleated from centrosome *c* is calculated by:

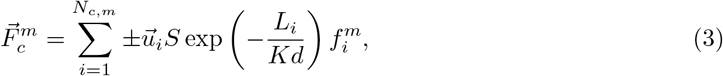

where the sign indicates the direction of the force (cortical dynein, dynein at spindle poles, and Eg5 are (−) while HSET is (+)). The exponential term accounts for increased drag-dependent force as the centrosome approaches a boundary, i.e. the cell cortex or opposing centrosome (Aponte-Rivera and Zia, 2016). Therefore, this term is dependent on the distance, *L_i_*, between the centrosome and the point of force application on microtubule *i*, and the distance, *d*, either between centrosomes or between the centrosome and the cell cortex. *K* is a constant scaling factor, and *S* is an additional motor-dependent scaling. For dynein-dependent forces, where only one microtubule is bound, *S* = 1. For HSET and Eg5-derived forces, where two antiparallel microtubules are interacting, *S* = *a*(1 + *O_i,j_*)*C*, where *a* is a constant that depends on the angle between interacting microtubules, *O_i,j_* is the overlap distance between interpolar microtubules, and *C* is a constant scaling factor to account for both active and passive crosslinking activity at antiparallel microtubule overlap regions (Mollinari et al., 2002; Peterman and Scholey, 2009; Shimamoto et al., 2015; Reinemann et al., 2018; Lamson et al., 2019; Edelmaier et al., 2020).

All motor-dependent and non-motor-dependent forces on all microtubules emanating from centrosome c are summed to determine the total force on the centrosome, and the position of centrosome *c* is updated as:

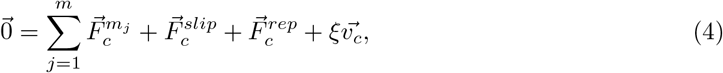

where 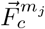 is the total force generated on centrosome *c* by motor *m_j_* (cortical dynein, HSET, Eg5, or spindle pole dynein), 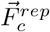 is the repulsive force on centrosome *c* when within a distance 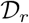 of another centrosome, 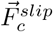 is the total slipping force on centrosome *c*, 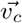 is the velocity of centrosome *c*, and *ξ* is a constant drag coefficient.

All parameters were optimized to match experimental results. Additionally, all parameters that were modified from our previous publication (Mercadante et al., 2021) were re-tested in simulations with two centrosomes and asymmetric dynein localization to confirm that appropriate bipolar spindle length and dynamics were maintained (Supplemental Figure S8).

### Modulating and Assessing Cortical Dynein Activity

To spatially and temporally regulate cortical dynein localization and activity within the model, we specify which microtubules are able to bind to cortical dynein with probability *P_d_cor__* based on the position of the microtubule end. When cortical dynein is uniformly distributed, we allow all microtubules to have an equal probability of binding to dynein and generating force. We remove cortical dynein by setting the probability *P_d_cor__* = 0.01, preventing most microtubules from binding and generating force. With dynein localized in a specific region as shown in Figure 4 A and D, we only allow microtubules whose end falls within the upper-right quadrant of the cell to bind to cortical dynein at a probability *P_d_cor__*. We allow all other microtubules throughout the cell to have a probability *P_d_cor__* = 0.01 under the assumption that cortical dynein is unlikely to be entirely absent from this region. When regions of dynein localization oscillate in time, we change the position requirement of microtubule ends to bind to cortical dynein, dependent on the period of dynein oscillations, *T*. Specifically microtubules in either the upper-right or lower-left quadrant bind to cortical dynein with probability *P_d_cor__* while all other microtubules have a small probability *P_d_cor__* = 0.01 of binding. Peaks in the traces defining the pairwise distances between centrosomes was determined by the MATLAB function “findpeaks” (MathWorks, 2007), with significant peaks defined as those having a prominence greater than 1 standard deviation of the average peak prominence.

### Computation and Code Availability

All computational modeling and model analysis was performed in MATLAB utilizing GPUs on a cluster. Thirty simulations of each set of parameters were run with three centrosomes for 30 minutes of mitosis unless otherwise specified. Code will be made available upon request.

### Cell Culture

Cell lines were maintained at 37°C with 5% CO_2_. hTERT-immortalized Retinal Pigment Epithelial (RPE) cells were obtained from and authenticated by ATCC. RPE cells expressing EGFP-tubulin were generated by viral transduction of L304-EGFP-Tubulin, a gift from Weiping Han (Addgene plasmid #64060; http://n2t.net/addgene:64060; RRID:Addgene_64060; Yang et al. (2013)). RPE cells expressing the tet-inducible PLK4, a gift from Neil Ganem (Boston University) and MDA-MB-231 cells, a gift from Catherine Whittington (Worcester Polytechnic Institute) were maintained in Dulbecco’s Modified Essential Medium (DMEM). RPE p53 deficient cells, a gift from Meng-Fu Bryan Tsou (Memorial Sloan Kettering Cancer Center) were maintained in DMEM F-12. Cell culture medium was supplemented with 10% Fetal Bovine Serum (FBS) and 1% Penicillin and Streptomycin. DNA stain (DAPI) was used to monitor and confirm absence of *Mycoplasma* contamination.

### Induction of centrosome amplification and Perturbation of Cortical Dynein Activity

Disruption of cortical localization of dynein was achieved through depletion of Afadin or LGN using lipid based transfection of 50 nM Horizon ON-TARGET plus pools of siRNA (Afadin target sequences: 5’-ugagaaaccucua guugua-3’, 5’-ccaaaugguuuacaagaau-3’, 5’-guuaagggcccaagacaua-3’, 5’-acuugagcggcaucgaaua-3’, LGN target sequences: 5’-gaacuaacagcacgacuua-3’, 5’-cuucagggaugcaguuaua-3’, 5’-acagugaaauucuugcuaa-3’, 5’-ugaaggguucuuugacuua-3’). Horizon non-targeting siRNA pool (siCtl) was used as a negative control for siRNA experiments (5’-ugguuuacaugucgacuaa-3’, 5’-ugguuuacauguuguguga-3’, 5’-ugguuuacauguuuucuga-3’, 5’-gguuuacauguuuuccua-3’). Knockdown efficiency was confirmed using Afadin (F:5’-gtgggacagcattaccgaca-3’, R:5’tcatcggcttcaccattcc-3’), LGN (F:5’-gtgaccacccgtctgtcg-3’, R:5’-ttcagcaacatttctcccgc-3’), and GAPDH (F:5’-ctagctggcccgatttctcc-3’, R:5’-cgcccaatacgaccaaatcaga-3’) specific primers. Inhibition of cortical dynein activity was achieved with exposure to 25 *μ*M of dynarrestin for 1 hour. Four hours post transfection with siRNA, RPE indPLK4 cells were induced to express PLK4 and amplify centrosome biogenesis by addition of 2 *μ*g/mL of Doxycycline four hours post transfection with siRNA. PLK4 induction was sustained for 48 hours until cell fixation. Alternatively, 24 hours after siRNA transfection, RPE p53-/-cells were treated with 1.5 *μ*g/mL Dihydrocytochalasin B (DCB) for 24 hours to induce cytokinesis failure and generate tetraploid cells with 4 centrosomes. Cells were washed out of DCB and cultured for an additional 24 hours prior to allowing tetraploid cells to progress to mitosis. These experimental timelines are summarized in Supplemental Figure S2. Where relevant, 20 *μ*M MG132 was added to the media for the final 30 minutes or 1hour of culture prior to fixation.

### Fluorescence Imaging and Analysis

Images were captured with a Zyla sCMOS (Oxford Instruments, Belfast, UK) camera mounted on a Nikon Ti-E microscope (Nikon, Tokyo, Japan). For live cell imaging of RPE cells expressing EGFP-tubulin, a 20x CFI Plan Fluor objective was used to capture images every 2.5 minutes for the duration of mitosis (Mercadante et al., 2019). Analysis of mitotic timing and cell fate was performed on at least 50 mitotic cells. Mitotic duration was quantified as the time between nuclear envelope breakdown (determined by loss of GFP exclusion from the nucleus) to anaphase B (determined by rapid elongation of the spindle). Phase contrast images were used to assess the number of progeny resulting from each division.

For analysis of cortex-localized NuMA, cells were rinsed briefly in phosphate buffered saline (PBS), then incubated in PHEM buffer with 0.3% Triton X-100 (TX-100) for 5 minutes. Cells were fixed in warmed 3.7% paraformaldehyde supplemented with 30 mM sucrose for 15 minutes at room temperature and then permeabilized in 0.1% TX-100 in PBS for 5 minutes. Cells were blocked in 3% PBS-BSA with 0.05% Tween-20 for 15 minutes and then incubated in primary antibody (NuMa: Abcam ab109262, Cambridge, UK; centrin-2: Sigma 04-1624) diluted in blocking buffer overnight at 4°C. Cells were washed briefly and incubated in secondary antibody diluted in TBS-BSA containing 0.2 *μ*mg/mL DAPI. NuMA distribution was qualitatively assessed with respect to centrosome positioning. For analysis of spindle morphology, cells were fixed in ice cold methanol for 10 minutes at −20°C. Blocking, primary (α-tubulin: Abcam ab18251, Cambridge, UK; Centrin: Millipore 04-1624, Burlington, MA) and secondary antibodies dilutions were prepared in TBS-BSA. DNA was detected with the addition of 0.2 *μ*g/mL DAPI to the secondary antibody dilution. Images of individual mitotic cells were captured using a 60× Plan Apo oil immersion objective and 0.3 *μ*m z-stacks through the depth of the cell. NuMA localization was assessed as described previously (Seldin et al., 2013). Clustered bipolar spindles were defined as those in which microtubules were organized into two spindle poles and in which one or both spindle poles contained 2 or more centrosomes positioned within 5 *μ*m of each other. Statistical analysis between two conditions was determined using a two-tailed student’s t-test. For multiple comparisons, a one-way ANOVA was performed and Dunnett’s test post-hoc was used for simultaneous comparison between each test condition and a control.

## Acknowledgments

Results in this paper were obtained in part using a high-performance computing system acquired through NSF MRI grant DMS-1227943 to WPI. This work was supported by NSF GRFP to DLM and NIH R01 GM140465-01 to ALM and SDO.

## Supplementary Material

A supplement to this article contains additional figures and information about the computational model.

## Notes

### Competing Interest Statement

The authors have declared no competing interest.

