## Supplemental Figures for "Cortical Dynein Drives Centrosome Clustering in Cells with Centrosome Amplification"

Table S1: Parameters. All parameter values without reference are approximated to match biological results.

| Parameter | Value | Description | Reference |
|---|---|---|---|
| **Microtubules** | | | |
| $v_g$ | 0.183 $\mu$ms$^{-1}$ | Microtubule growth velocity (+ ends) | Piehl and Cassimeris (2003), Wu et al. (2011) |
| $v_s$ | 0.3 $\mu$ms$^{-1}$ | Microtubule shrinking velocity (+ ends) | Wu et al. (2011) |
| $v_b$ | 0.057 $\mu ms^{-1}$ | Microtubule shrinking velocity (+ ends) bound to cortical dynein | Laan et al. (2012) |
| $k_1$ | 0.167 s$^{-1}$ | Rescue frequency | Wu et al. (2011), Komarova et al. (2002) |
| $\kappa$ | 10 pN$\mu$m$^2$ | Bending rigidity | Laan et al. (2012), Li and Jiang (2017), Kikumotot et al. (2006) |
| $f_{stall}$ | 5 pN | Stall force of microtubules | Ma et al. (2013) |
| $MT_{nuc}$ | 2 s$^{-1}$ | Microtubule nucleation rate per centrosome | Piehl et al. (2004) |
| $\theta$ | $10\pi/180$ | Slipping microtubule angle change | |
| **Motor Proteins** | | | |
| *Dynein* | | | |
| $f_{0,d}$ | 3.6 pN | Stall force of dynein | Elshenawy et al. (2019) |
| $v_{0,d}$ | 0.86 $\mu$ms$^{-1}$ | Walking velocity of dynein | Urnavicius et al. (2018), Elshenawy et al. (2019) |
| $P_{d_{cor}}$ | 0.5 | Probability of binding to cortical dynein | |
| $P_{d_{sp}}$ | 0.3 | Probability of binding to spindle pole dynein | |
| $\mathcal{D}_{d_{cor}}$ | $4v_g(dt)$ $\mu$m | Distance required for binding to cortical dynein | |
| $\mathcal{D}_{d_{sp}}$ | 1 $\mu$m | Distance required for binding to dynein at spindle poles | |
| *Kinesin-5 (Eg5)* | | | |
| $f_{0,Eg5}$ | 1.5 pN | Stall force of Eg5 | Shimamoto et al. (2015) |
| $v_{0,Eg5}$ | 0.2 $\mu$ms$^{-1}$ | Walking velocity of Eg5 | Li et al. (2019) |
| $P_E$ | 0.5 | Probability of binding to Eg5 | |
| *Kinesin-14 (HSET)* | | | |
| $f_{0,HSET}$ | 1.1 pN | Stall force of HSET | Roostalu et al. (2018) |
| $v_{0,HSET}$ | 0.2 $\mu$ms$^{-1}$ | Walking velocity of HSET | Li et al. (2019) |
| $P_H$ | 0.7 | Probability of binding to HSET | |
| $\mathcal{D}_{Eg5,HSET}$ | $v_g dt$ $\mu$m | Distance required for binding to Eg5 or HSET | |
| **Other** | | | |
| $r$ | 15 $\mu$m | Radius of the cell | |
| $c_r$ | 0.3 $\mu$m | Radius of a centrosome | |
| $\mathcal{D}_r$ | 2 $\mu$m | Distance for repulsive forces | |
| $K$ | 0.35 | Microtubule length-dependent scaling factor | |
| $C$ | 0.01 | Antiparallel crosslinking scaling factor | |
| $s$ | 0.075 $\mu$m$^{-1}$ | Scaling for catastrophe frequency | |
| $R$ | 1 $\mu$m | Scaling for repulsive forces | |
| $\mu$ | 0.7 pNs$\mu$m$^{-2}$ | Viscosity of the cytoplasm | Luby-Phelps et al. (1993) |
| $\xi$ | 41.2 pNs | Drag coefficient | |

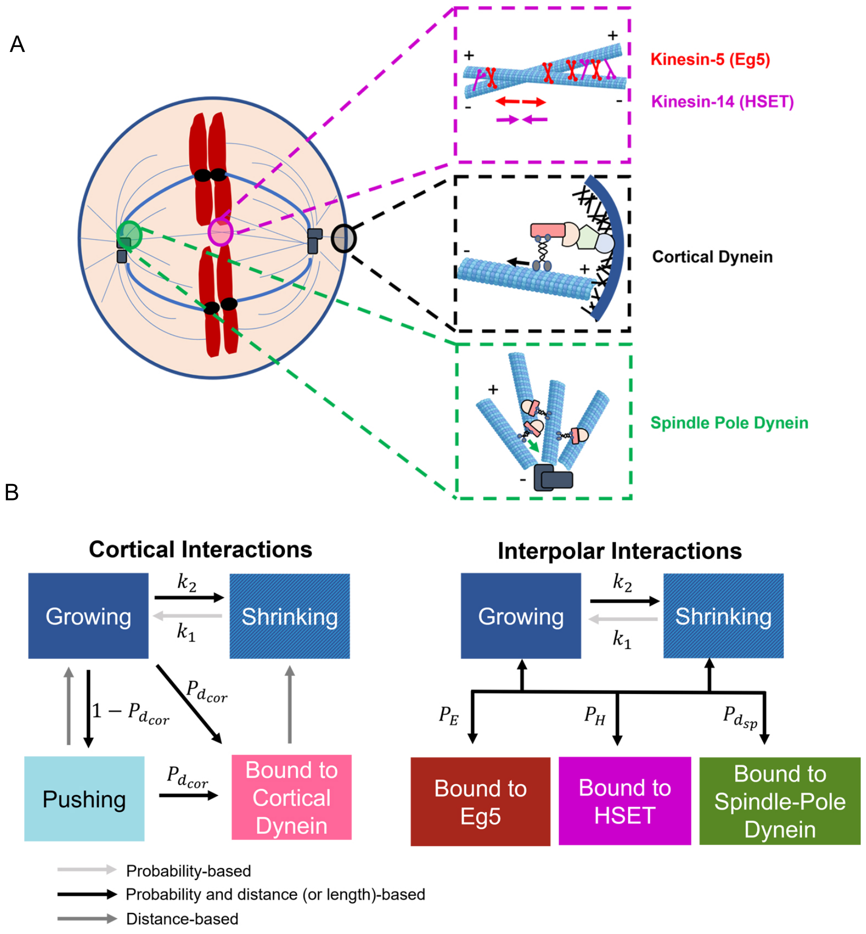


Figure S1: Microtubule dynamics and motor-derived forces are dependent on probabilities of binding and/or distances between microtubule and motor. (A) Schematic of the mitotic spindle and microtubule-motor protein interactions. Arrows depict the directional movement of the motor along the microtubule. (B) Schematic representation of stochastic microtubule states (left) and microtubule-motor protein interactions (right).

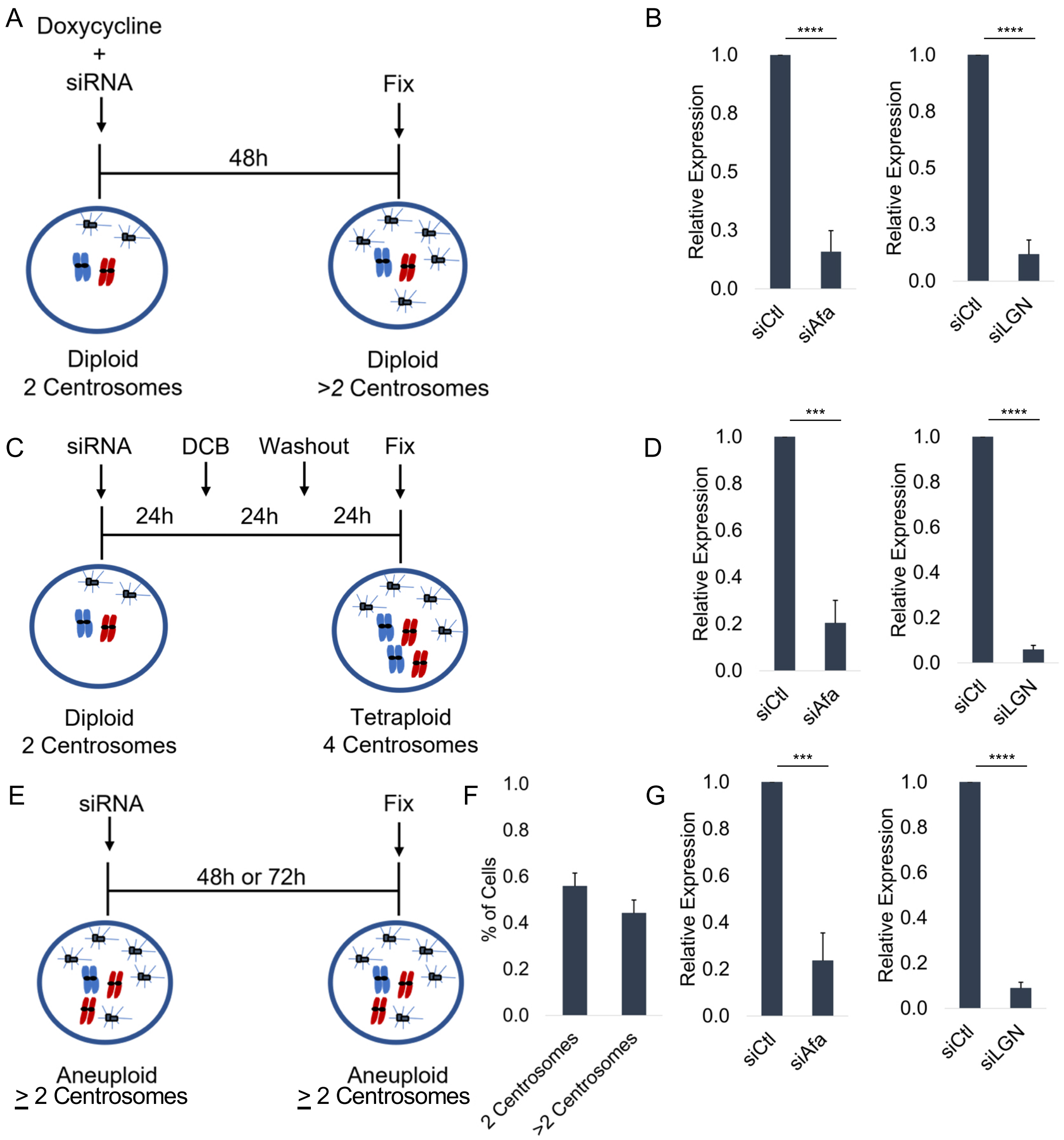


Figure S2: Experimental timeline for induction of centrosome amplification and disruption of cortical dynein localization. (A,C) Depiction of the experimental timelines to induce siRNA-based depletion of Afadin or LGN in cells concurrently induced to have centrosome amplification through either overexpression of PLK4 (A) or dihydrocytochalasin B (DCB)-induced cytokinesis failure (C). (B,D) Relative expression of Afadin (left) or LGN (right) following indicated duration of siRNA-mediated depletion of Afadin and LGN. (E) Quantification of centrosome number in mitotic MDA-MB-231 cancer cells. (F) Relative expression of Afadin (left) or LGN (right) after 72 h or 48 h, respectively, of siRNA-mediated depletion. Significance determined by student's t-test; ***$p < 0.001$, ****$p < 0.0001$.

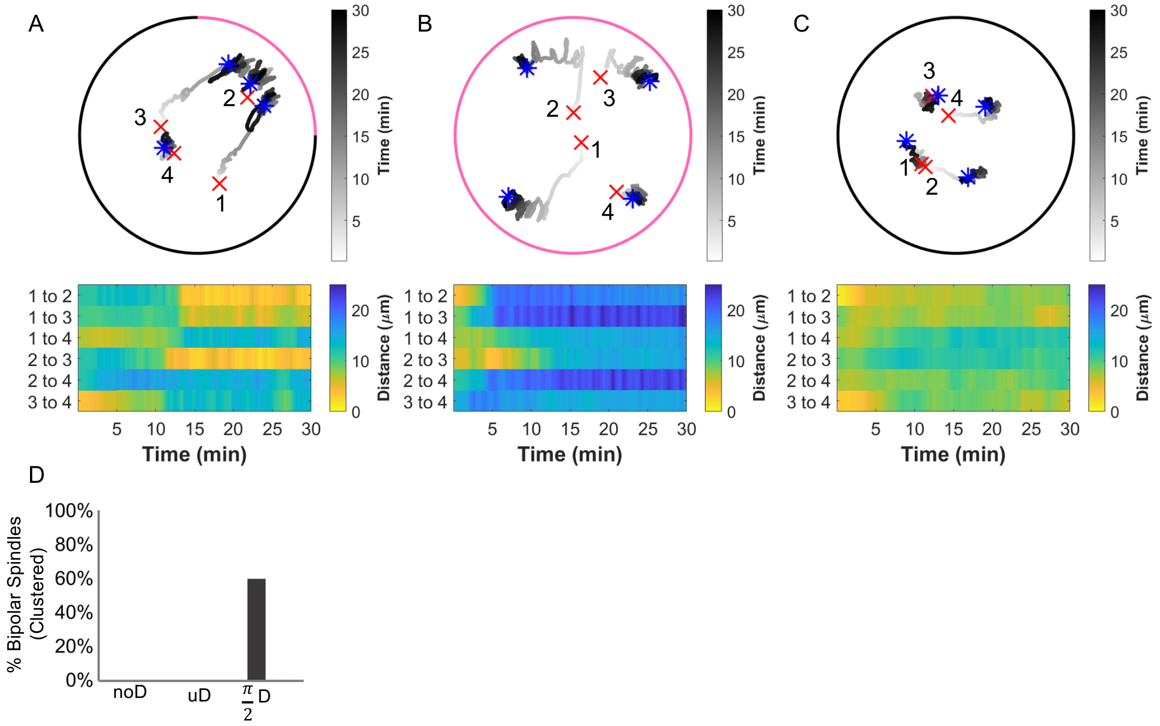


Figure S3: Centrosome movement and clustering is similar with 3 or 4 centrosomes. (A-C) Top: Trace of centrosome movement over time through the duration of a simulation where a red 'x' indicates initial centrosome position and a blue '*' indicates centrosome position at 30 minutes. Numbers mark individual centrosomes, grayscale indicates time, and pink on the cell boundary indicates the region of high dynein localization (where $P_{d_{cor}} = 0.5$; elsewhere $P_{d_{cor}} = 0.01$). Bottom: Heat map representing the pairwise distances between all centrosome pairs indicated in the corresponding traces. (D) The percent of simulations that achieve bipolar spindles (centrosome clustering) when cortical dynein is absent (noD), distributed uniformly across the cell boundary (uD), or enriched in the angular region from 0 to $\pi/2$ (($\pi/2$)D). All simulations are for t=30 minutes and data in (D) represents 10 simulations for conditions shown in (A-C).

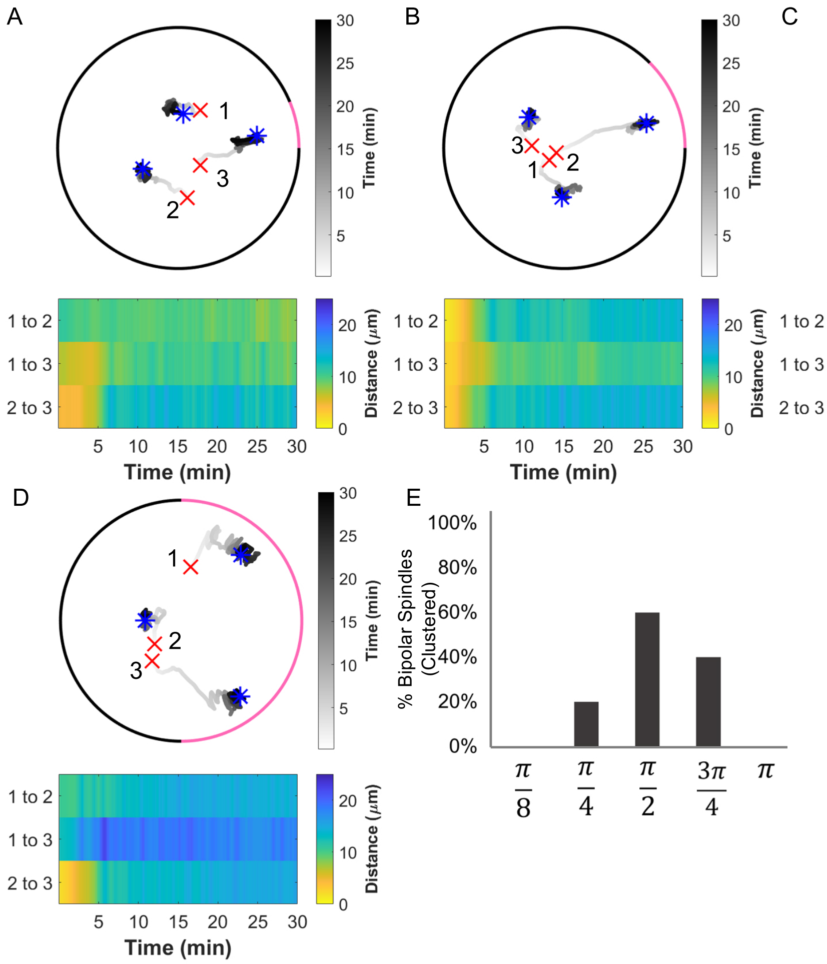


Figure S4: Centrosome clustering is sensitive to the size of the region of cortical dynein activity. (A-D) Top: Trace of centrosome movement over time through the duration of a simulation where a red 'x' indicates initial centrosome position, a blue '*' indicates centrosome position at 30 minutes. Numbers mark individual centrosomes, grayscale indicates time, and pink on the cell boundary indicates the region of high dynein activity (where $P_{d_{cor}} = 0.5$; elsewhere $P_{d_{cor}} = 0.01$). Dynein is enriched in a region of size $\pi/8$, $\pi/4$, $3\pi/4$, and $\pi$ in panels A, B, C, and D, respectively. Bottom: Heat map representing the pairwise distances between all centrosome pairs indicated in the corresponding traces. (E) The percent of simulations that achieve centrosome clustering (pairwise distance$\leq$5 $\mu$m) when cortical dynein activity is enriched in a region of size $\frac{\pi}{8}$, $\frac{\pi}{4}$, $\frac{\pi}{2}$ (condition represented in Figure 4 A,B), $\frac{3\pi}{4}$, and $\pi$. Data in E represents 5 simulations for each condition.

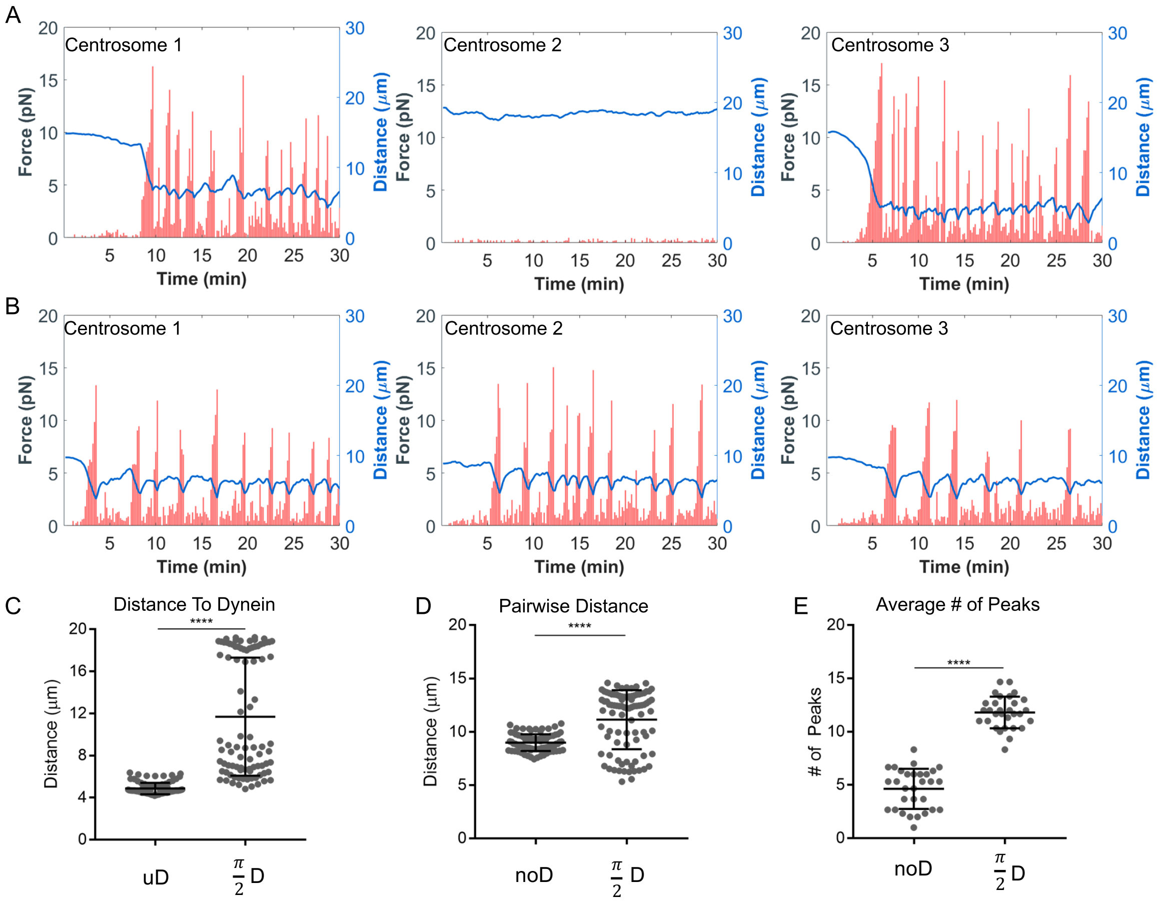


Figure S5: Centrosome movement is driven by cortical dynein activity. (A,B) Plots depicting the magnitude of cortical dynein-derived forces over time (red bars, scale on left axis) and the distance between each centrosome and the midpoint of dynein enrichment (blue trace, scale on right axis) from the simulation with dynein enriched in the angular region from 0 to $\pi/2$ in A and uniform dynein in B (corresponding to simulations in Figure 4 B and D, respectively). (C) Distance between each centrosome and the region of dynein enrichment. (D) Distance between centrosome pairs. For simulations with uniform dynein, the distance is calculated as the minimum distance to the cell boundary. (E) Average number of peaks in cortical-dynein-derived force in simulations. In (C-E), noD, uD, and $(\pi/2)$D correspond to the cases of no dynein, uniformly distributed dynein, or dynein restricted to the angular region from 0 to $\pi/2$. Each dot represents a single centrosome's position (C), centrosome pair (D), or simulation (E). Quantification performed on 30 simulations from each condition. Significance determined by one-way ANOVA with Tukey's test for multiple comparisons; $^{*}p < 0.05$, $^{****}p < 0.0001$.

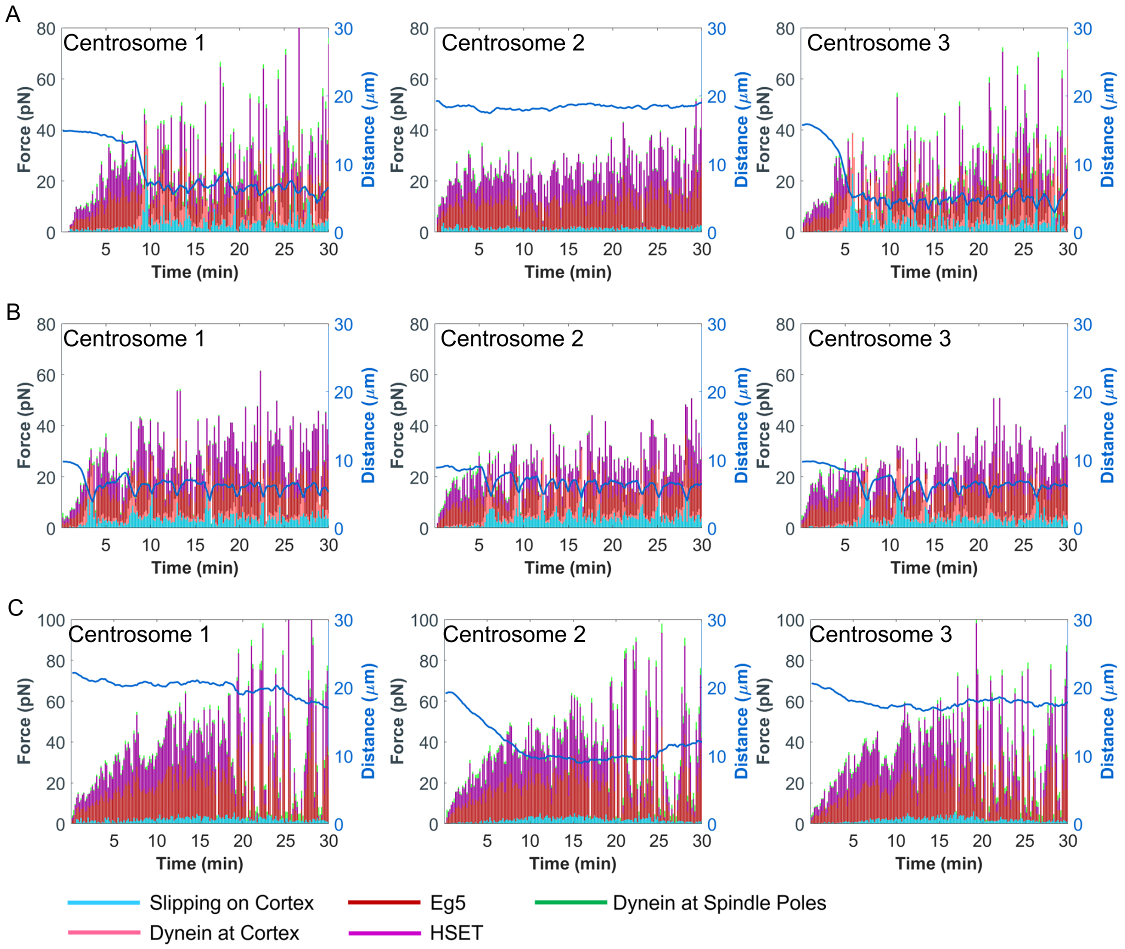


Figure S6: All motor and non-motor derived forces are dynamic over time. Plots depicting the magnitude of each force (scale on left axis) on each centrosome, and the corresponding centrosome position relative to the cell cortex (blue trace, scale on right axis) over time. Plots correspond to simulations as follows: (A) Figure 4B, (B) Figure 4D, and (C) Figure 4E.

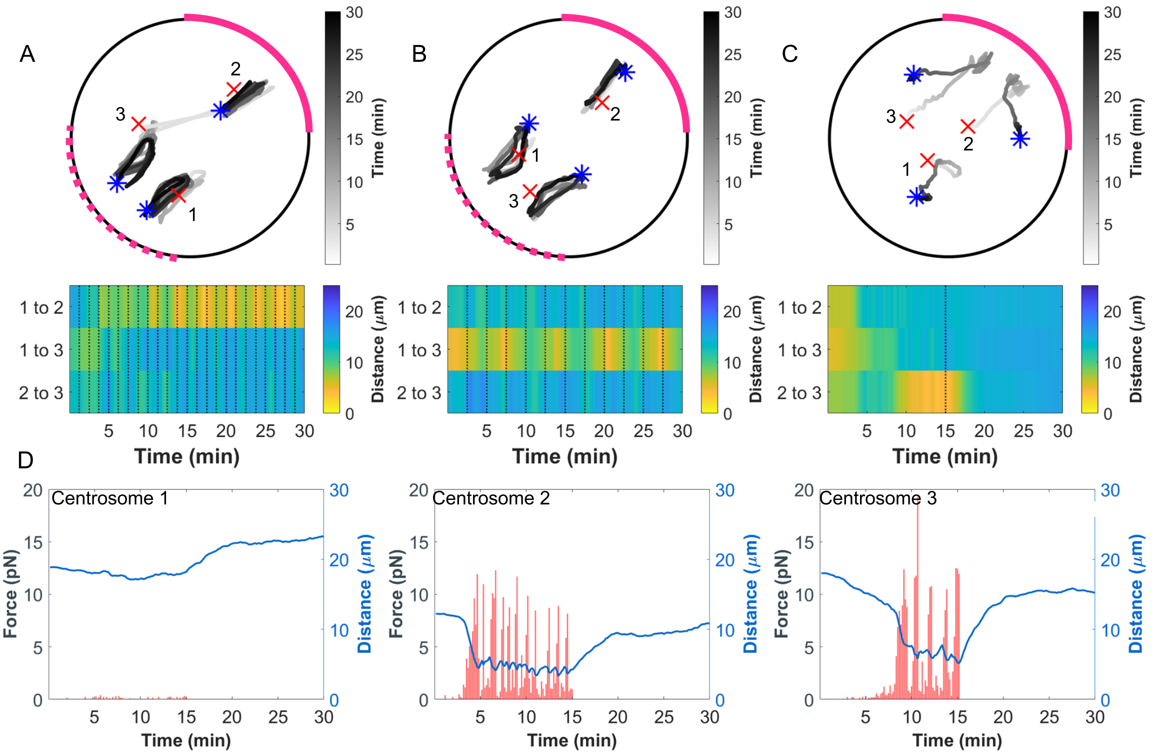


Figure S7: Asymmetric and dynamic cortical dynein localization aids centrosome clustering. Top: Traces of centrosome movement over time from a simulation with enriched region of cortical dynein localization (with $P_{d_{cor}} = 0.5$) oscillating between the upper-right (solid pink arc, 0 to $\pi/2$) and lower-left quadrant (dashed pink arc, $-\pi$ to $-\pi/2$) of the cell with a period of $T = 1.67$ min (A) and $T = 2.33$ min (B). In (C), enrichment is only up to $T = 15$ min and then there is no dynein enrichment for $T \geq 15$ ($P_{d_{cor}} = 0.01$ everywhere on the cortex). Initial centrosome position indicated by red a 'x' and final centrosome position indicated by a blue '*'. Color bar is time. Bottom: Heat map representing the pairwise distances between all centrosome pairs from the simulation represented above. Black dotted lines indicate a timepoint when cortical dynein localization is redistributed. Color bar is distance. (D) Plots depicting the magnitude of cortical dynein-derived forces over time (red bars, scale on left axis) and the distance (blue traces, scale on right axis) between each centrosome and the midpoint of dynein localization from the simulation shown in (C).

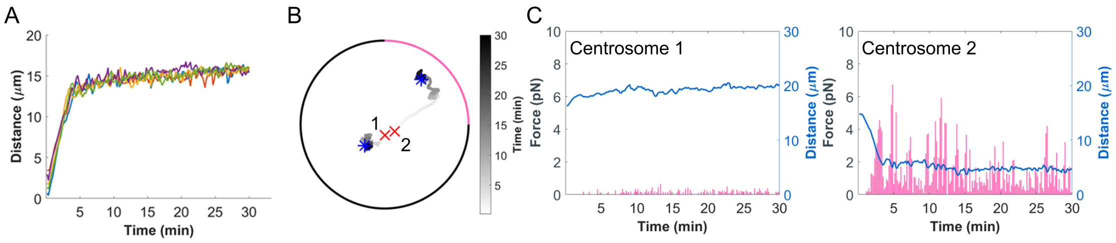


Figure S8: Spindle length and dynein-derived forces are dynamic over time in cells with two centrosomes. (A) Traces of bipolar spindle length from 5 simulations with asymmetric dynein localization. (B) Traces of centrosome movement over time from a simulation with asymmetric dynein localization. Colorbar is time. (C) Plots depicting the magnitude of cortical dynein-derived forces (pink bars, scale on left axis) over time and the distance (blue trace, shown on right axis) between each centrosome and the midpoint of dynein localization from the simulation shown in (B).
